# Conformational Dynamics of NSP11 Peptide of SARS-CoV-2 Under Membrane Mimetics and Different Solvent Conditions

**DOI:** 10.1101/2020.10.07.330068

**Authors:** Kundlik Gadhave, Prateek Kumar, Ankur Kumar, Taniya Bhardwaj, Neha Garg, Rajanish Giri

**Affiliations:** School of Basic Sciences, Indian Institute of Technology Mandi, Kamand, Himachal Pradesh, 175005, India; Department of Medicinal Chemistry, Faculty of Ayurveda, Institute of Medical Sciences, Banaras Hindu University, Varanasi, Uttar Pradesh, 221005, India

**Keywords:** COVID-19, SARS-CoV-2, nsp11, CD Spectroscopy, Intrinsically disordered proteins

## Abstract

The intrinsically disordered proteins/regions (IDPs/IDPRs) are known to be responsible for multiple cellular processes and are associated with many chronic diseases. In viruses, the existence of a disordered proteome is also proven and is related to its conformational dynamics inside the host. The SARS-CoV-2 has a large proteome, in which, structure and functions of many proteins are not known yet, along with nsp11. In this study, we have performed extensive experimentation on nsp11. Our results based on the CD spectroscopy gives characteristic disordered spectrum for IDPs. Further, we investigated the conformational behavior of nsp11 in the presence of membrane mimetic environment, α-helix inducer, and natural osmolyte. In the presence of negatively charged and neutral liposomes, nsp11 remains disordered. However, with SDS micelle, it adopted an α-helical conformation, suggesting the helical propensity of nsp11. Finally, we again confirmed the IDP behavior of nsp11 using MD simulations. In future, this conformational dynamic study could help to clarify its functional importance in SARS-CoV-2 infection.

## 1. Introduction

The existing pandemic caused by severe acute respiratory syndrome coronavirus 2 (SARS-CoV-2) is an enveloped RNA virus. Its genome organization shows the presence of a ~30 kbp positive-sense single-stranded RNA (+ssRNA) buried inside a layer of viral proteins involved in protecting and shaping the virion particle [1–3]. Once inside the host cell, genomic RNA is translated into three different types of viral proteins: structural, accessory, and non-structural proteins (nsps). Structural proteins (spike, envelope, membrane, and nucleocapsid) together form the outer cover of virion [1,4]. Accessory proteins aid in virus survival by evading the host immune system, inducing cell death, and modulating viral pathogenicity. The third category of coronavirus proteins comprises of the nsps which are cleaved from two translated polyproteins (pp1a and pp1ab) by two proteases (PLpro and 3CLpro) [4–6]. These virus-encoded nsps serves as an essential component for replication of genome and transcriptional activity. In fact, several nsps are involved throughout initial and intermediate phases of the viral life cycle [7]. The recently emerged SARS-CoV-2 has a short protein, nsp11, of only 13 amino acids [5]. Generally, depending on the CoVs species, the nsp11 comprises 13–23 residues [7]. The nsp11 protein is the cleavage product of pp1a polyprotein by 3CLpro/Mpro protease at the nsp10/11 junction (putative cleavage site is- nsp10 PMLQ|SADA nsp11) [8]. Interestingly, voluntary ribosomal frameshifting occurs in coronaviruses in replicase gene (−1 frameshift) giving rise to rest of the nsp12-16 proteins of longer polyprotein 1ab [7].

In our recent study, we analysed the propensity of intrinsic disorder in SARS-CoV-2 proteins and correlated it with its evolutionary closer human SARS and Bat SARS-like coronaviruses [9]. The multiple sequence alignment of SARS-CoV-2 nsp11 with SARS-CoV and Bat SARS-like coronavirus represents the difference in protein sequence at 5^th^ and 6^th^ positions where glutamine and serine are observed to substitute serine and threonine residues, respectively [9]. As an uncharacterised protein of potential function, published literature on nsp11 protein of murine hepatitis virus shows cleavage mutants nsp10/nsp11 are not replication viable [10]. On contrarily, cleavage mutants nsp10-nsp11/nsp12 of avian coronavirus infectious bronchitis virus are dispensable for viral replication [6]. Sun *et al.* reported that the overexpression of nsp11 of porcine reproductive and respiratory syndrome virus (PRRSV) induced a strong suppression of interferon (IFN) production [11].

The release and fate of nsp11 protein in CoV-infected cells have not been recognised so far. Moreover, its structural, biophysical, and functional information in any coronavirus has still not been elucidated which is vital to understand its biology and pathogenesis from a structural point of view. In this study, using Far-UV CD spectroscopy, we identified SARS-CoV-2 nsp11 as an intrinsically disordered protein (IDP). IDPs are unstructured proteins that lack structural constraints forming an ensemble of structures [12,13]. In contrast to classical structure-function-paradigm, IDPs works either through structure-independent interactions or by virtue of gaining a secondary structure (called as MoRFs) such as α-helix on interaction with their physiological partner [14,15].

Several proteins from organisms of all three kingdoms of life have been studied to perform structure-independent functions [16,17]. Viruses are no less behind and also have multiple flexible proteins [18–20]. The sequence-dependent characteristic of IDPs enables them to have an exceptionally large interactome [16,17]. We, therefore, as an essential step in direction of understanding nsp11 protein and its interacting ability, employed experimental as well as computational methods to analyse its ‘gain-of-structure’ capacity. We used various model systems such as DOPC, DOPS, SDS micelles, TMAO, and TFE to mimic the biological environments like hydrophobic-hydrophilic interface, and protein-lipids interaction. Collectively, our findings revealed the unstructured nature of SARS-CoV-2 nsp11 protein along with membrane-mediated effects of liposomes on its structure. The comprehensive structural investigation may help to deduce the function and structure-function relationship of nsp11 protein in SARS-CoV-2. As nsp11 is present at the site where ribosomal frameshift occurs, the role of disordered nature of nsp11 therein needs to be elucidated.

## 2. Material and Methods

### 2.1. Material

The 2,2,2-trifluoroethanol (TFE) and Sodium dodecyl sulfate (SDS), and trimethylamine N-oxide (TMAO) were procured from Sigma-Aldrich (USA). The lipids 1,2-dioleoyl-sn-glycero-3-phosphocholine (DOPC) and 1,2-dioleoyl-sn-glycero-3-phospho-L-serine (DOPS) were obtained from Avanti Polar Lipids (Alabaster, Alabama, U.S.A.) and prepared using the lipid extruder provided by the same manufacturer. nsp11 peptide (NH2-SADAQSFLNGFAV-COOH; UniProt ID: P0DTC1 (residues 4393-4405)) was procured from *Gene* Script with > 80 % purity. The peptide was dissolved in 20 mM phosphate buffer (pH 7.4) containing 50 mM NaCl to prepare a stock solution (3 mg/ml).

### 2.2. Liposome preparation

The chloroform from lipids (DOPC/DOPS) solution was removed using rotatory evaporators. Afterward, overnight incubation was done in a desiccator to remove any traces of chloroform. The dried lipid was then hydrated with 20 mM sodium phosphate buffer (pH 7.4) and further lipid suspension (DOPC: 29 mg/ml; DOPS: 20 mg/ml) was processed five times by freeze (liquid N2)/thaw (60° C water bath)/vortex cycles. Lipid suspension was then subjected to extrusion to prepare large unilamellar vesicles (LUVs) using 0.1 μm pore diameter polycarbonate membrane. For the extrusion process, we used Avanti mini-extruder (Avanti) and followed the manufacturer’s protocol. The prepared LUVs were stored at 4°C and used within 5 days.

### 2.3. Dynamic light scattering (DLS)

The hydrodynamic radius/size of the prepared LUVs was measured by using DLS (Zetasizer Nano S from Malvern Instruments Ltd., UK). The prepared LUVs were diluted in water (1:100 ratio of LUV and water) before measurements. The observed size of the DOPC and DOPS was 102 nm and 124 nm respectively.

### 2.4. Circular dichroism (CD) spectroscopy

The far UV-CD spectrum of nsp11 protein was recorded in J-1500 spectrophotometer (Jasco) at 25°C. We used 1 mm optical path length quartz cuvette to record spectra in various concentrations of SDS, LUVs, TFE, and varying temperature condition. The scan range spanned 190–240 nm, with a 0.5 nm data interval. The 0.1 mm optical path length cuvette was used to record spectra in the case of TMAO. 20 mM phosphate buffer (pH 7.4, 50 mM NaCl) was used to prepare the nsp11 sample (100 μM) for a spectral scan. 500 μM of nsp11 were used in the case of TMAO. The CD spectrum was recorded as an average of 3 consecutive scans. Each spectrum was fitted by Savitsky-Golay smoothing (15 points of smoothing window and second polynomial order). The secondary structure contents (α-helix, β-sheet, disordered, and turn) of the samples were calculated by deconvolution of CD spectroscopic data in DICHROWEB webserver (CONTIN algorithm with a reference set 7) [21].

### 2.5. Structure modelling and molecular dynamics (MD) simulation

The long timescale computer simulations have been very useful to comprehend the atomic-level dynamics of a peptide like nsp11 of SARS-CoV-2. There is very less information available about its intrinsic properties. Thus, to gain atomic insight on dynamics of nsp11, we have performed all-atom molecular dynamics simulations. Firstly, a 3D model was constructed for the 13 amino acids long peptide using PEP-FOLD peptide structure prediction server [22]. It applies a coarse-grained (CG) forcefield and performs up to 200ns simulation runs which generate upto 200 models. These models are clustered using OPEP (Optimized Potential for Efficient structure Prediction) and then build an energy minimized structure as used previously [23,24]. The resultant model was then prepared using Chimera by addition of missing hydrogens and proper assignment of bond orders. We used Charmm36 (march 2019) forcefield in Gromacs (v2018.8), where simulation setup was built by placing the protein structure in a cubic box along with TIP3P water model [25]. After solvation, the system was charge neutralized with counterions. To attain an energy minimized simulation system, the steepest descent method was used until the system was converged within 1000 kJ/mol. Further, the equilibration of system was done to optimize solvent in the environment. Using NVT and NPT ensembles within periodic boundary conditions for 100ps each, the system was equilibrated. The average temperature at 300K and pressure at 1 bar were maintained using Nose-Hoover and Parrinello-Rahman coupling methods during simulation. All bond-related constraints were solved using SHAKE algorithm. The final production run was performed for 500ns in our high performing cluster at IIT Mandi. After analyzing the trajectory, the last frame (NSP11LF) was chosen for further studies in different conditions.

Next, the structural conformations of NSP11 using NSP11LF were analyzed in two different conditions of solvents: TFE (8M) and SDS (60 molecules) mixed with water in separate simulation runs. The topology of TFE and SDS molecules were generated by PRODRG2 webserver [26]. By using Gromacs 54a7 forcefield parameters and addition of TFE and SDS molecules into the system, the simulations were performed for 200ns and 500ns for TFE and SDS based systems, respectively.

Further, Replica Exchange MD was performed for 100 ns of last frame (NSP11LF) using our previous protocol in Desmond simulation package [27]. Here, we have chosen 8 temperatures for replica-exchange based on experiments. The temperatures were calculated by linear mode in Desmond in 8 replicas viz. 298K (25°C), 307.29K (34.29°C), 316.57K (43.57°C), 325.86K (52.86°C), 335.14K (62.14°C), 344.43K (71.43°C), 353.71K (80.71°C), and 363K (90°C).

All trajectory analysis, calculations were performed using Chimera, Maestro and Gromacs commands for calculating the root mean square deviation (RMSD), root mean square fluctuation (RMSF), and radius of gyration (Rg) for protein structure compactness for C-α atoms.

## 3. Result and Discussion

### 3.1. SARS-CoV-2 nsp11 is intrinsically disordered under physiological buffer conditions

The Far-UV CD spectrum of nsp11 protein shows a signature minima at 200 nm (**Figure 1A**), suggest the spectrum of typical disordered type protein.

**Figure 1:**
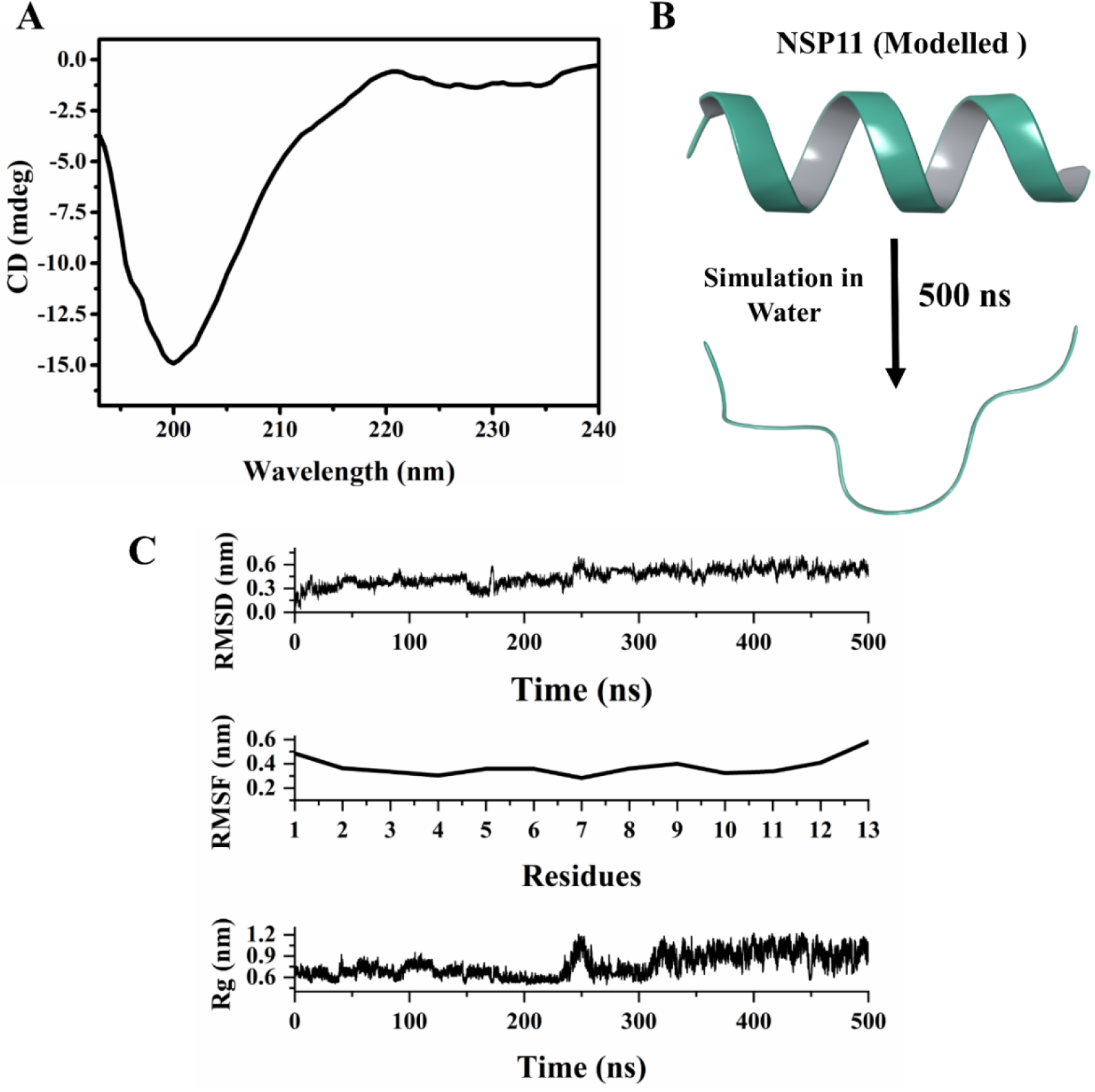
Secondary structure analysis and MD simulation of nsp11. (A) Far-UV CD spectra of nsp11 represent characteristic disordered spectra (B) Structure of nsp11 obtained from molecular modelling in the PEP-FOLD server and further MD simulation performed at 500 ns in water. (C) RMSD, RMSF, and Rg analysis in reference to the initial structure of MD trajectory.

The structural insight into the nsp11 was further investigated through all-atom MD simulations up to 500 ns. The structure of nsp11 was entirely in helical conformation after modelling by PEP-FOLD webserver. After preparing the structure for its proper symmetry and hydrogen addition, the structure was simulated in presence of SPC water model for 500 ns. As observed in simulations, the structure loses its helicity during simulation period (**Figure 1B**). In terms of mean fluctuations and deviations at the atomic scale, high values for RMSD, RMSF, and Rg were detected (**Figure 1C**). After 250 ns, the RMSD values were increased from approximately 0.25 nm to 0.6 nm and remained stable till 500 ns. Similarly, the Rg increased 0.65 nm to 1.1 nm in radius of gyration that suggests lower compactness as compared to the initial structure. Also, RMSF is showing higher fluctuation at the terminal residues between 0.4 to 0.6 nm as compared to the middle segment. Higher fluctuation and deviation may be due to the presence of disordered structure in nsp11. The disordered proteins are highly dynamic where the simulation and CD spectra revealed nsp11 as disordered protein. However, the prediction and homology modelling suggest that the nsp11 also has some helical structure indicating that it has an intrinsic ability to form helix when environmental condition demands.

### 3.2. Disordered nsp11 shows an upsurge in α-helical content in TFE

The lipids/membrane mimetics, natural osmolytes, and organic solvents (α-helix inducer) are generally used to explore the folding propensities of disordered proteins [28]. As nsp11 is a disordered system it is wise to use TFE, SDS, Lipids, and osmolytes to investigate the fold propensity and structural transition from disordered to ordered one. The TFE is well-characterised to initiate folding and induce secondary structures in protein chains. It indirectly stabilizes the intra H-bonds of an α-helix by destabilizing or weakening the H-bonds between CO and NH groups of protein backbone with water [29]. With increasing TFE concentration it is evident that the negative ellipticity at 200 nm becomes lesser and negative ellipticities at 208 nm and 222 nm become towards higher value (**Figure 2A, B**). The change in the ellipticity at the corresponding wavelength indicates that nsp11 acquires an α-helical structure in presence of TFE. Additionally, the transition from disorder to order structure is accompanied by the formation of an intermediate structure near 20 % of TFE (**Figure 2B**) representing a partially folded conformation. **Table 1** represents the increase in α-helical and decrease in disordered contents in varying TFE concentration.

**Figure 2:**
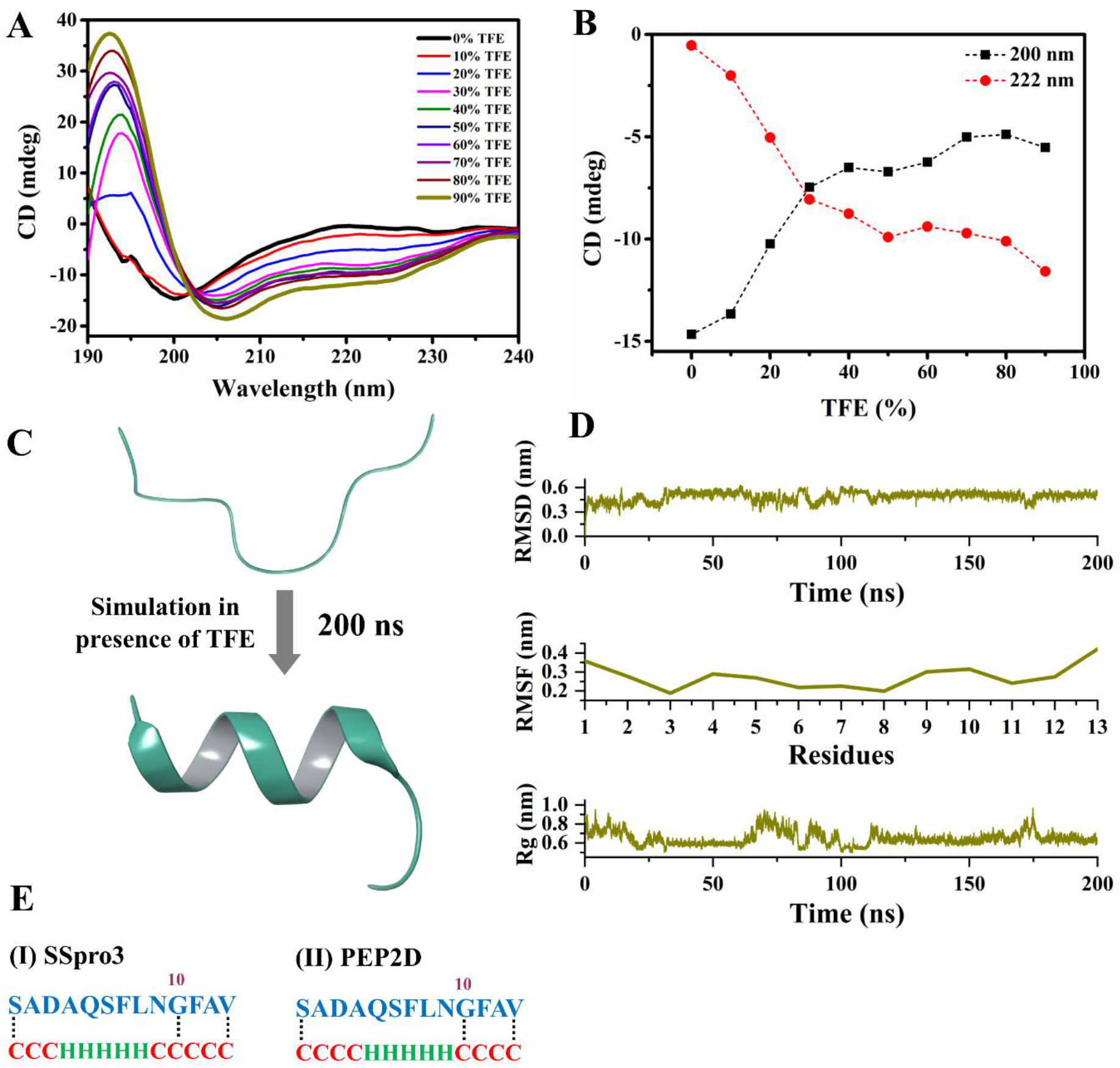
Secondary structure analysis of nsp11 in TFE. (A) Effect of TFE (0-90%) concentration on conformation of disordered nsp11 protein monitored by Far-UV CD. Higher TFE concentration induces α-helix in nsp11. Plot (B) signifies the change of ellipticity at 200 nm (black squares with dashed line) and 222 nm (red circles with dashed line) as the function of an increasing TFE concentration. (C) Disordered to ordered transition of nsp11 in presence of TFE at 200 ns. 3D structure of nsp11 in presence of TFE at 200 ns showing a helical structure in the middle segment (4AQSFLNGF11). (D) The fluctuating trend in RMSD, RMSF, Rg in reference to the initial structure of MD trajectory depicts the occurring changes for nsp11 in TFE. (E) Secondary structure perdition for nsp11 using (I) SSPro3 and (II) PEP2D (H-Helix (green) and C-Coil (red)) to predict helix promoting region in nsp11.

**Table 1:** Estimation of secondary structure content of nsp11 in presence SDS and TFE from DICHROWEB CD analysis.

|  |  | Secondary structure content (%) |  |  |  |
| --- | --- | --- | --- | --- | --- |
| | | α-helix | β-sheet | Turn | Disordered |
| SDS | 0 mM | 8.85% | 11.75% | 12.4% | 66.95% |
|  | 0.5 mM | 8.25% | 14.95% | 12.05% | 64.75% |
|  | 100 mM | 98.85% | 1.15% | 0% | 0% |
| TFE | 10% | 8.05% | 18.95% | 11.75% | 61.25% |
|  | 50% | 87.4% | 12.55% | 0% | 0% |
|  | 90% | 98.65% | 1.35% | 0% | 0% |

Further, simulations in presence of TFE produce a remarkable change in the structure of nsp11 which is consistent with the result obtained from Far-UV CD. A disorder to order (α-helix) transition is seen at residues 4AQSFLNGF11 (**Figure 2C**) which shows stable RMSF. But both the terminals are showing higher fluctuation due to disordered terminal. RMSD and Rg show lesser fluctuation due to higher structured content (**Figure 2D**). Further, to know α-helix promoting regions of the nsp11, we performed sequence-based secondary structure prediction by SSpro3 (from SCRATCH protein predictor) [30] and PEP2D [31] servers (**Figure 2E**). SSpro3 server predicted residues 4^th^-8^th^ as α-helix (38.46%) and residues 1^st^-3^rd^ and 9^th^-13^th^ as random coil (61.54%) whereas the PEP2D server predicted residues 1-4^th^ and 10-13^th^ as a random coil (61.54%) and residues 5-9^th^ as α-helix (38.46%) (**Figure 2E)**. This is in correlation with MD results. The structural changes in TFE may arise due to disordered proteins lacking a stable three-dimensional conformation at the physiological condition and easily adopts a secondary structure in the presence of a favourable environment or suitable binding partner [13].

### 3.3. nsp11 adopts α-helix structure in SDS but remain unaffected in negatively charged and neutral lipids

The membrane inside cells are very dynamic and therefore, artificially prepared models can be used to study protein-membrane or protein-protein interactions [32]. Here, LUVs with neutral DOPC and anionic DOPS were prepared and used for CD experiments. Several IDPs are reported to bind efficiently to the DOPC and DOPS lipids and this interaction is accompanied by an enhanced level of their α-helical structure [23,28,33]. In the case of nsp11, the presence of DOPS and DOPC does not show any change in secondary structure conformation (**Figure 3E and 3F**). The nsp11 has one negatively charged residue and does not contain positively charged residues that may have a role in its interaction with lipids. Analogous to lipid membranes, the SDS micelles used to mimic the interface between hydrophobic and hydrophilic environments such as the plasma membrane and the cytosol [34]. SDS is a denaturing detergent with folding inducing properties that form micelles (CMC is 1.99 mM in 50 mM phosphate buffer [35]) and interacts with hydrophobic parts of proteins [36]. At lower SDS concentration (below CMC) where SDS is in the monomeric forms do not shows any structural change and the shape of the spectra remains similar as of nsp11 in the absence of SDS. Whereas at 5mM (above CMC) the shape of the spectrum changes and exhibits a shift in negative ellipticity from 200 nm to 203 nm. At higher concentrations (25, 50, and 100 mM) the CD spectra of nsp11 exhibited two minima at 206 and 222 nm suggesting the structural transition from the disordered to ordered conformation (**Figure 3A, B**). Further, CD deconvolution confirms the reduction in disordered structure content and an increase in ordered structure (α-helix) content (**Table 1**) in nsp11 above CMC.

**Figure 3:**
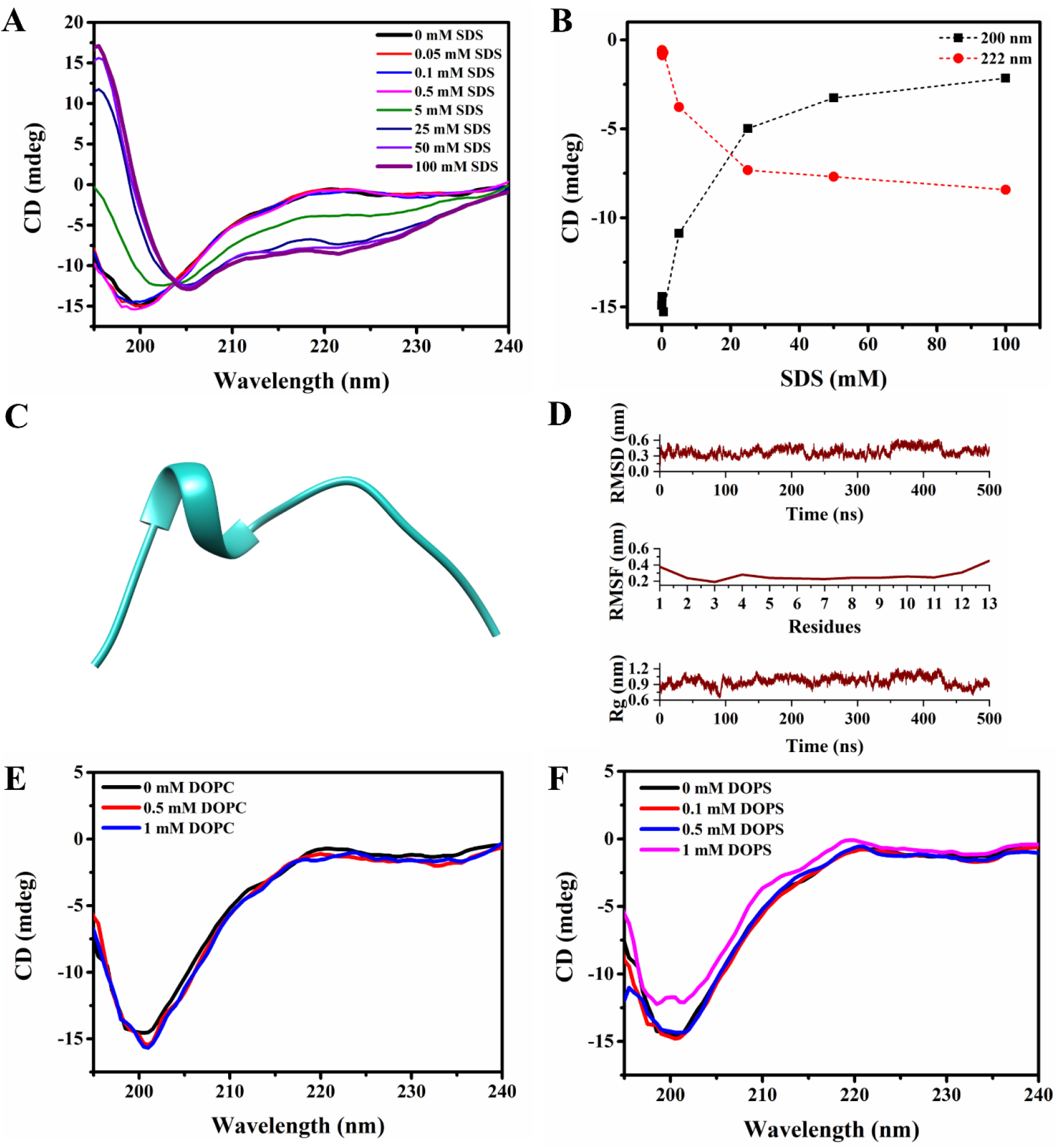
Effect of Membrane-Mimetic Environment on nsp11 secondary structure. (A) Far-UV CD spectra of nsp11 in presence of SDS. Increasing concentration of SDS induces helical content in nsp11. (B) representation of ellipticity at 200 nm (black squares with dashed line) and 222 nm (red circles with dashed line) as the function of SDS concentration. (C) MD simulation trajectory of nsp11 in presence of SDS. The last frame (at 500 ns) is showing propensity of the helix (4AQSF7). (D) RMSD, RMSF, and Rg analysis in reference to the initial structure in presence of SDS molecules. **(E)** Far-UV CD spectra of nsp11 measured in presence of neutral lipid (DOPC) and (F) anionic lipid DOPS. Both lipids are unable to induce any structural change in nsp11.

Like CD spectroscopy, MD simulations of nsp11 in presence of SDS molecules is also showing some noticeable change in structure as shown in **Figure 3C**. A significant proportion of helix was formed by four residues 4AQSF7 at the last frame (500 ns) simulation which contributes to nearly 30% of ordered structure. A little fluctuation was observed in the case of RMSD and Rg may be due to the disordered terminal region which shows high RMSF throughout the simulation (**Figure 3D**). The RMSF of residues at the middle segment is stable that suggests the gain of the ordered structure during simulation in SDS. Collectively, the in-depth insight of structural transition from disordered to ordered one suggests the possible interaction of nsp11 with membrane mimetics which may explain the host specificity to membrane disruption implicated in the viral pathogenesis.

### 3.4. Osmolytes show no effect on secondary structure of nsp11

In the cell, various classes of organic osmolytes are present and may have several roles therein [28,37]. Thus, natural osmolytes such as TMAO were used on structural properties of nsp11 protein. The osmophobic effect of TMAO force thermodynamically unstable proteins or disordered proteins to fold and regain high functional activity [28]. In a disordered state, the peptide backbone is largely exposed to its surrounding, as an osmolyte, TMAO exerts a solvophobic thermodynamic force which raises the free energy of disordered state preferentially shifting the equilibrium towards the folded form. This osmophobic effect drives the folding of natively unstructured proteins [38,39]. In the case of nsp11, TMAO is not exerting any effect on its secondary structure. The shape of the far-UV CD spectra remains the same irrespective of increased TMAO concentration (up to 2 M). Only a slight shift in signature minimum was observed (shift from 200nm to 203 nm) at 3.25 M TMAO but the shape of the spectra retains its disordered type protein (**Figure 4**).

**Figure 4:**
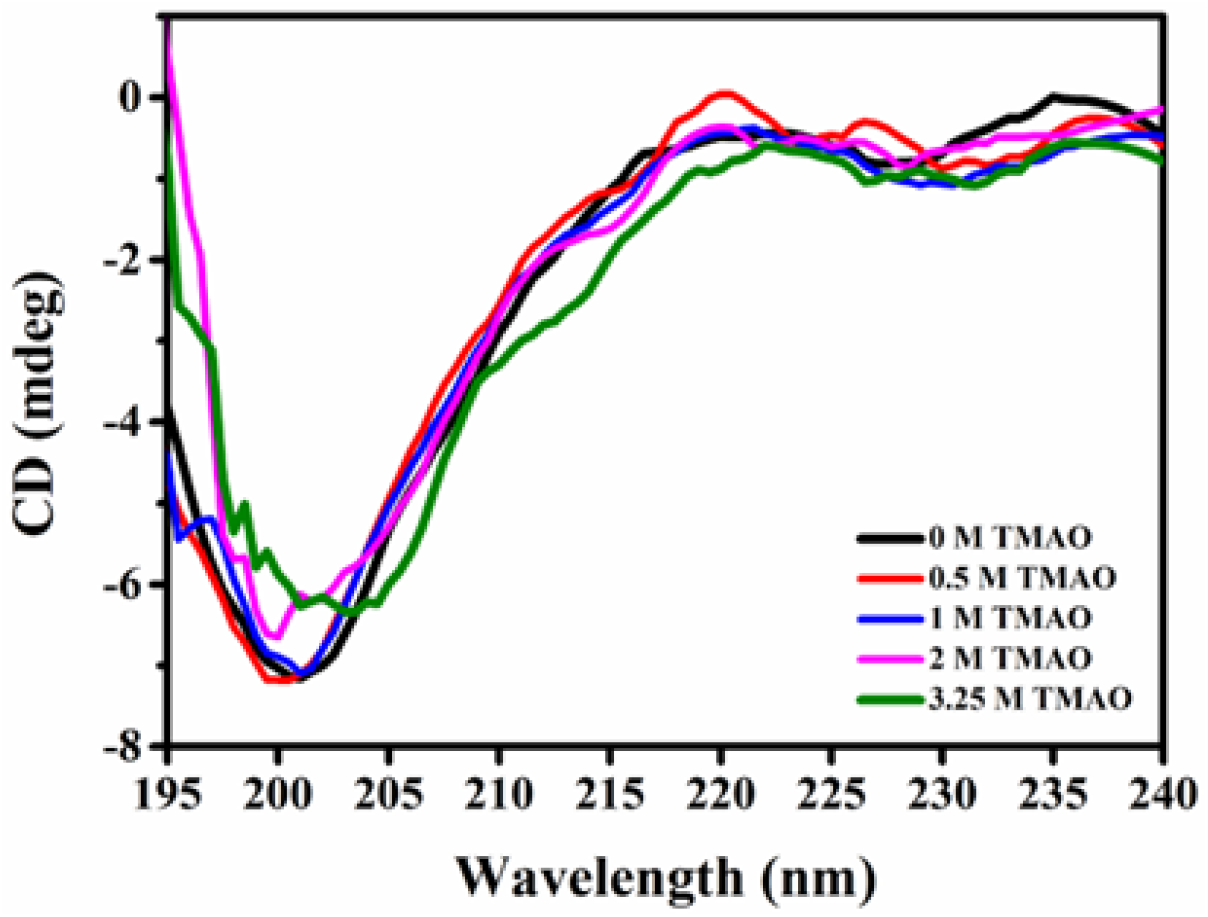
Far-UV CD spectra of nsp11 in presence of TMAO. Irrespective of the TMAO concentration the shape of CD spectra represents the typical disordered type protein suggesting TMAO unable to induce secondary structure change in nsp11.

### 3.5. Temperature-induced structural change in nsp11 appeared by contraction in structure

Further, to investigate the temperature-induced conformational changes in nsp11 we acquired CD spectra at 10 °C to 90 °C with 5 °C interval as shown in **Figure 5A**. At lower temperatures, the far-UV CD spectrum is typical of an unfolded protein. As the temperature rises, the signature minima remain nearly at 200 nm, but the shape of the spectrum shows some changes at higher temperature. With increasing temperature, the ellipticity (negative) corresponding to signature minima at 200 nm becomes lower and higher at 222 nm (**Figure 5B**).

**Figure 5:**
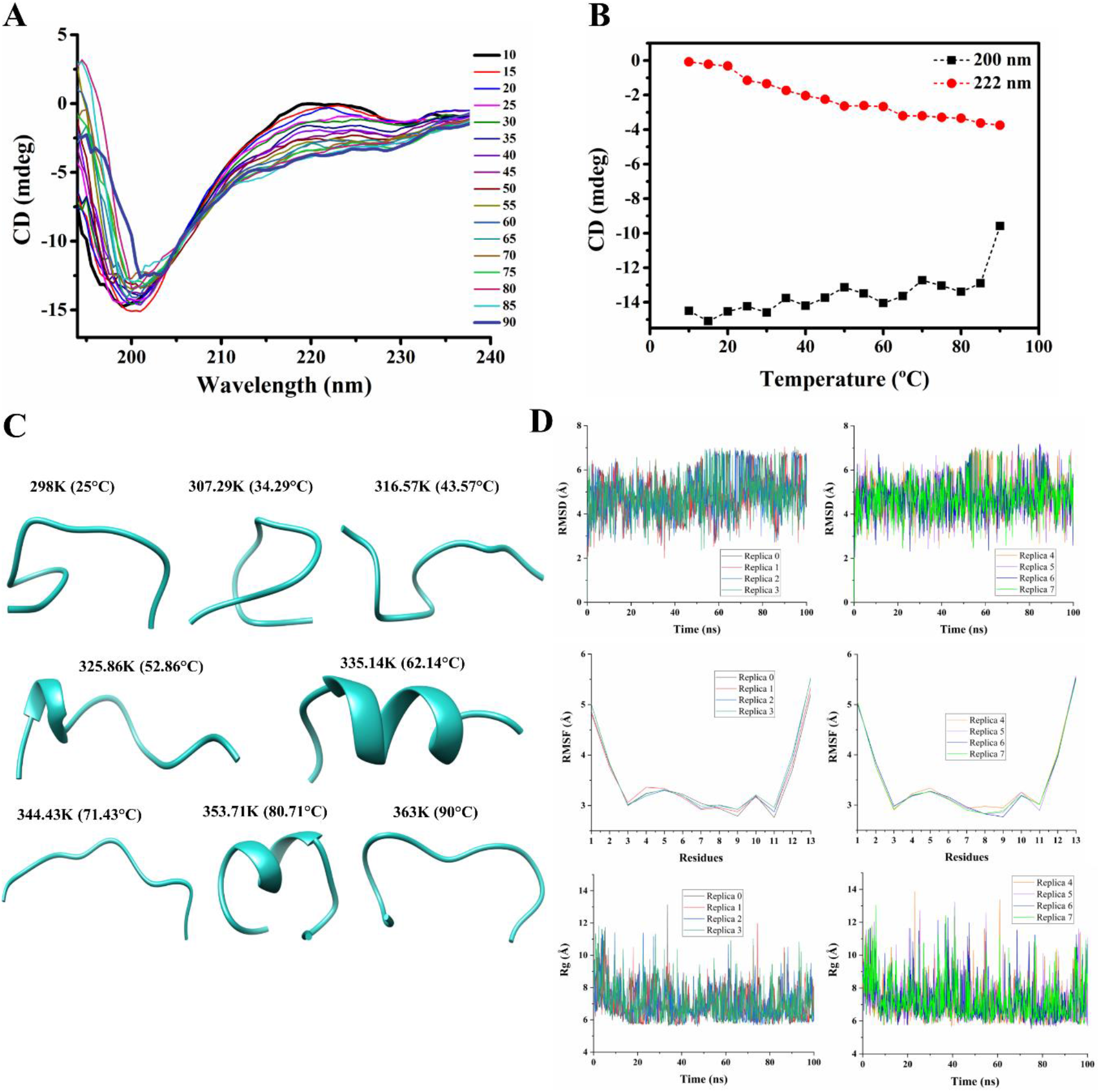
Effect of temperature on secondary structure of nsp11 protein. (A) Thermal stability of nsp11 secondary structure monitored by far-UV CD spectra at temperatures range from 10°C to 90°C with 5°C interval. (B) Analysis of CD spectra obtained in fig A. black line represents change of ellipticity at 198 nm and red line represents change of ellipticity at 222 nm. (C) Last frame of each replica (0-7) of REMD simulation with increasing temperature. (D) RMSD, RMSF, and Rg analysis in reference to the initial structure of each trajectory.

The change in the shape of the spectra at higher temperatures, indicating temperature-induced partial folding or partial formation of ordered secondary structure. This type of change is in line with the other disordered proteins [23,33,40]. The structural changes in IDPs at higher temperature is the common phenomenon which attended by the contraction of the conformational ensemble [41]. The contraction of structure occur in nsp11 may be due to change in secondary or tertiary structure [41]. Moreover, reports suggested that the folding in the disordered proteins at higher temperatures may occur due to enhancement in the strength of hydrophobic interaction by elevated temperature [28]. The hydrophobicity value of an nsp11 protein was calculated according to ProtParam [42] analysis shows the grand average of hydropathicity (GRAVY) score is 0.500 represents its hydrophobic nature (increasing positive score indicates a greater hydrophobicity). However, Nettels *et al* reported that that the contraction of disordered proteins is independent of the hydrophobicity of the protein, and therefore may not be due to enhanced hydrophobic interactions [43]. Some reports also suggested that the structural changes at higher temperatures are independent of long-range interactions and are occurred with the contraction of the conformational ensemble [41].

Further to gain insights into temperature-induced contraction in nsp11 we employed MD simulation where temperature-based replica exchange MD was performed to see the conformational change. Total 8 replicas were run and allow them to exchange conformations during MD simulation (**Figure 5C**). The structural change is not consistent during the simulation where nsp11 exists mostly in a disordered state. But at the higher temperature (52.86°C, 62.14°C, and 80.71°C) nsp11 gain helical structure. The helical structure gain is restricted mostly at the middle segment of the protein whereas both terminal residues remain disordered. It is also evident from RMSF where residues fluctuation for 3^rd^ to 11^th^ is stable but the fluctuation is very high for residues 1-2 and 12-13 (**Figure 5D**). Additionally, a more fluctuating peak of RMSD and Rg (**Figure 5D**) is suggesting that the nsp11 is mostly disordered irrespective of temperature rise. Overall, increasing temperature endorsed substantial changes in the shape of CD spectra that suggest a partial change in secondary structure conformation and this change may be due to partial helical structure gain at higher temperatures revealed by MD simulation.

## 4. Conclusion

This study accentuates the structural conformation and dynamics of SARS-CoV-2 nsp11 protein. Our study signifies that the nsp11 is an intrinsically disordered protein. The model conditions mimicking the effect of the membrane field (TFE) is effective to induce helical structure in nsp11. It shows the high specificity towards the SDS micelles, compared to neutral and anionic lipids, and readily undergo a conformational transition (disorder to order) in the presence of SDS micelle. Importantly, the consequent structural transformations and dynamics at different membrane environments could be dependent on the type of solvent used. Thus, our study emphasizes that nsp11 may play a functional role in the host cytosolic membrane affinity/interaction.

## Future perspectives

In the wake of ongoing pandemic, understanding the biology and structure-function paradigm is essential to understand the pathogenesis of the virus. Especially, the identification of the role of such proteins which results in important events like ribosomal frameshift needs to be identified. In the last two decades, it has been established that proteins with no proper three-dimensional structure have a great amount of significance in most cellular processes. Herein, the identified intrinsic disordered nature of the SARS-CoV-2 nsp11 protein will surely help to investigate its role in the ribosomal frameshifting which can give some promising targets for identifying the contagious nature of SARS-CoV-2 virus.

## Abbreviations

IDPs: intrinsically disordered proteins
IDPRs: IDP regions
SARS-CoV-2: severe acute respiratory syndrome coronavirus 2
nsps: non-structural proteins
TFE: 2,2,2-trifluoroethanol
SDS: Sodium dodecyl sulfate
TMAO: trimethylamine N-oxide
DOPC: 1,2-dioleoyl-sn-glycero-3-phosphocholine
DOPS: 1,2-dioleoyl-sn-glycero-3-phospho-L-serine
LUVs: large unilamellar vesicles
CD: circular dichroism
MD: molecular dynamics
RMSD: root mean square deviation
RMSF: root mean square fluctuation
Rg: radius of gyration
CMC: critical micelle concentration

## Author Contributions

RG and NG: Conception, design, review, and writing of the manuscript. KG and AK performed experiments and PK performed the computational study. KG, AK, PK, TB analyzed data and wrote the manuscript. All authors have read and approved the manuscript.

## Acknowledgments

Authors are thankful to Indian Institute of Technology Mandi (BioX, AMRC center, and HPC facility) for all the facilities. KG is supported by fellowship component of the grant from Science and Engineering Research Board (SERB), India (Grant Number: CRG/2019/005603 to RG). AK was supported by a fellowship from the Ministry of Human Resource Development (MHRD). TB is grateful to the Department of Science and Technology (DST) for Inspire Fellowship. RG is thankful to IYBA Award (Grant Number: BT/11/IYBA/2018/06) by Department of Biotechnology (DBT), India.

## UniProt accession ID

P0DTC1

(The nsp11 protein sequence were retrieved from SARS-CoV-2 (UniProt ID: P0DTC1), in that residues 4393-4405 is the sequence for nsp11).

